# TRAFIKK: systematic prediction and mechanistic interpretation of anticancer drug synergies

**DOI:** 10.64898/2026.05.08.723755

**Authors:** Marco Fariñas, Viviam Bermúdez, Eirini Tsirvouli, Kristine Lippestad, John Zobolas, Tero Aittokallio, Kaisa Lehti, Åsmund Flobak

**Affiliations:** Department of Chemistry and Biomedical Science, Norwegian University of Science and Technology (NTNU), Trondheim, Norway; Department of Clinical and Molecular Medicine, Norwegian University of Science and Technology (NTNU), Trondheim, Norway; Department of Biology, Norwegian University of Science and Technology (NTNU), Trondheim, Norway; Akerhus Clinical Research Center (ACR), Division of Research and Innovation, Akerhus University Hospital, Lørenskog, Norway; Division of Economy and Finance, Akerhus University Hospital, Lørenskog, Norway; Department of Cancer Genetics, Institute for Cancer Research, Oslo University Hospital (OUH), Oslo, Norway; Oslo Centre for Biostatistics and Epidemiology (OCBE), University of Oslo, Oslo, Norway; Institute for Molecular Medicine Finland (FIMM), HiLIFE, University of Helsinki, Helsinki, Finland; Department of Microbiology, Tumor and Cell Biology, Karolinska Institute (KI), Stockholm, Sweden; The Cancer Clinic, St. Olavs Hospital, Trondheim, Norway; Department of Biotechnology and Nanomedicine, Sintef Industry, Trondheim, Norway

## Abstract

Effective drug combination therapies can improve cancer treatment, yet the mechanistic basis of drug synergy remains poorly understood. Most computational approaches prioritize predictive accuracy over molecular mechanistic interpretability, providing hence limited insights into how synergistic effects emerge across signalling contexts. We developed Trafikk, a molecular-signalling network-based framework that simulates drug perturbations in cell line-specific computational models to mirror functional outcomes in experimental combination screens. Across two independent large-scale datasets, Trafikk identified synergistic combinations with >77% recall. Functional response predictions revealed both conserved and context-dependent mechanisms. While AKT-MEK co-inhibition consistently disrupted coordinated survival and apoptotic signalling in 742 cell lines, PI3K-BCL2 synergy arose through distinct death programs shaped by cell-context-specific network constraints. Trafikk combines predictive performance with mechanistic interpretability, capturing how and why drug synergy emerges across cellular contexts. Source code, installation instructions and usage tutorial are freely available at https://github.com/druglogics/trafikk.

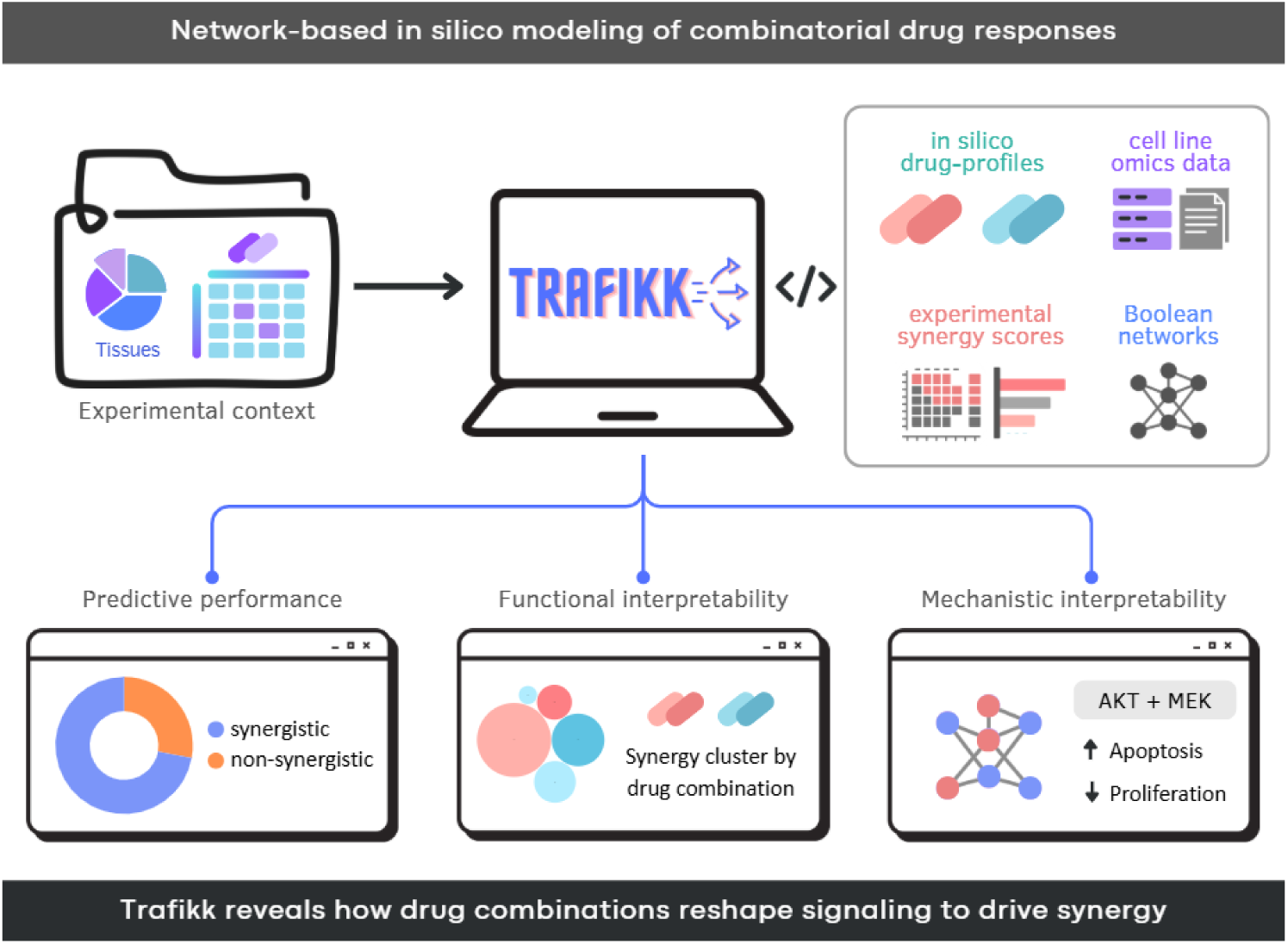

## Introduction

Combinatorial drug therapies have been suggested as a means to overcome adaptive and selective anti-cancer drug resistance. Discovery of effective combinations has, however, been challenged by yet poorly understood context-dependence, making one-size-fits-all discoveries rare and the experimental scale required for their discovery demanding. Large pan-cancer drug screens have confirmed that synergistic effects confer biological selectivity, with responses varying markedly across cellular backgrounds due to differences in mutations, pathway activities, and other context-dependent factors (1–3). While these studies highlight the necessity of finding effective drug combinations, they also expose the practical limitations of experimental screening alone, motivating the development of computational methods to prioritize promising drug pairs for testing their efficacy in pre-clinical cell models. Synergistic drug responses can be highly context-specific, reflecting that cancer arises from the dysregulation of interconnected signalling pathways rather than from isolated molecular interactions (4). These pathways form complex networks that integrate genetic, epigenetic, and environmental cues to regulate cell-fate decisions, such as proliferation and apoptosis. As a result, drug perturbations propagating through these networks rarely act in isolation, and are shaped by feedback mechanisms, which may not be fully captured by experimental screening alone (4, 5).

Advances in drug discovery frameworks have substantially improved the prediction of drug synergy by integrating chemical features, drug-target information, and cellular molecular profiles, enabling scalable and biologically informed modelling of nonlinear drug interactions (6, 7). However, although these approaches incorporate contextual molecular features to improve prediction accuracy, many operate as black boxes and provide limited insight into the mechanistic basis of synergistic responses (8). Network-based approaches offer a natural framework to address this limitation by explicitly modelling signalling interactions and their downstream consequences. In this context, Boolean modelling has been applied to simulate drug combination effects in cancer cells (9, 10) and other diseases (11, 12). Although Boolean modelling relies on discrete state representations, it provides a qualitative and mechanistic description of regulatory network dynamics that is well suited to capturing context-dependent signalling responses (6, 7).

Importantly, signal propagation analyses have shown that Boolean models can move beyond static outcomes, enabling a more nuanced characterization of how perturbations spread through Boolean networks and shape system-level responses (13, 14). In this context, we recently developed BooLEVARD, a computational framework that quantifies signal propagation in Boolean models (15). While Boolean network-based approaches have been applied to drug response modelling and combination analysis (16–18), existing methods typically focus on predictive performance and qualitative mechanistic exploration. To our knowledge, no framework has systematically integrated quantitative signal propagation, predictive assessment of drug synergy, and mechanistic interpretability, leaving an unmet need for frameworks that bridge mechanistic modelling with actionable drug prioritisation.

To address these limitations, we developed Trafikk, a computational framework to simulate drug perturbations in biological condition-specific Boolean models with signal-propagation analysis; the framework enables systematic generation of functional response profiles for single drugs and drug combinations. These profiles support network-level analyses that go beyond synergy classification, allowing identification and mechanistic interpretation of synergistic responses in terms of redistributed signalling and pathway engagement. Applied to two independent large-scale drug screens (1, 2), Trafikk robustly recapitulated experimentally observed synergistic responses across multiple cellular contexts and revealed that similar phenotypic outcomes can arise from distinct mechanistic signalling programs. This was exemplified with cases where single drugs act dominantly, as well as cases where non-redundant interactions between drugs give rise to broader combinatorial effects.

## Results

### Trafikk delivers a robust predictive performance in independent drug screens

To first evaluate whether the computational pipeline could reproduce experimentally observed drug synergy responses *in silico*, we assessed the performance of each model calibrated by all cell lines by comparing predictions to binarized Bliss synergy scores in two independent drug-screen datasets. The Sanger-2024 dataset was used for in-depth analyses across 757 cell lines due to broader model coverage of drug targets and combinations.

On the Sanger-2024 dataset, the overall performance of Trafikk across 16,460 cell line–drug combination comparisons reached a mean accuracy of 70.4% and a balanced accuracy of 66.7%, with both recall and precision at 77.8% (**Fig 1A**). Across 522 evaluable cell lines, the median AUC-ROC was 0.67 (mean 0.67 ± 0.14) and the median AUC-PR was 0.78 (mean 0.76 ± 0.15) (**Fig 1B**). Consistent performance was observed across cancer types (**Supplementary Figs S2** and **S3**).

**Fig 1.**
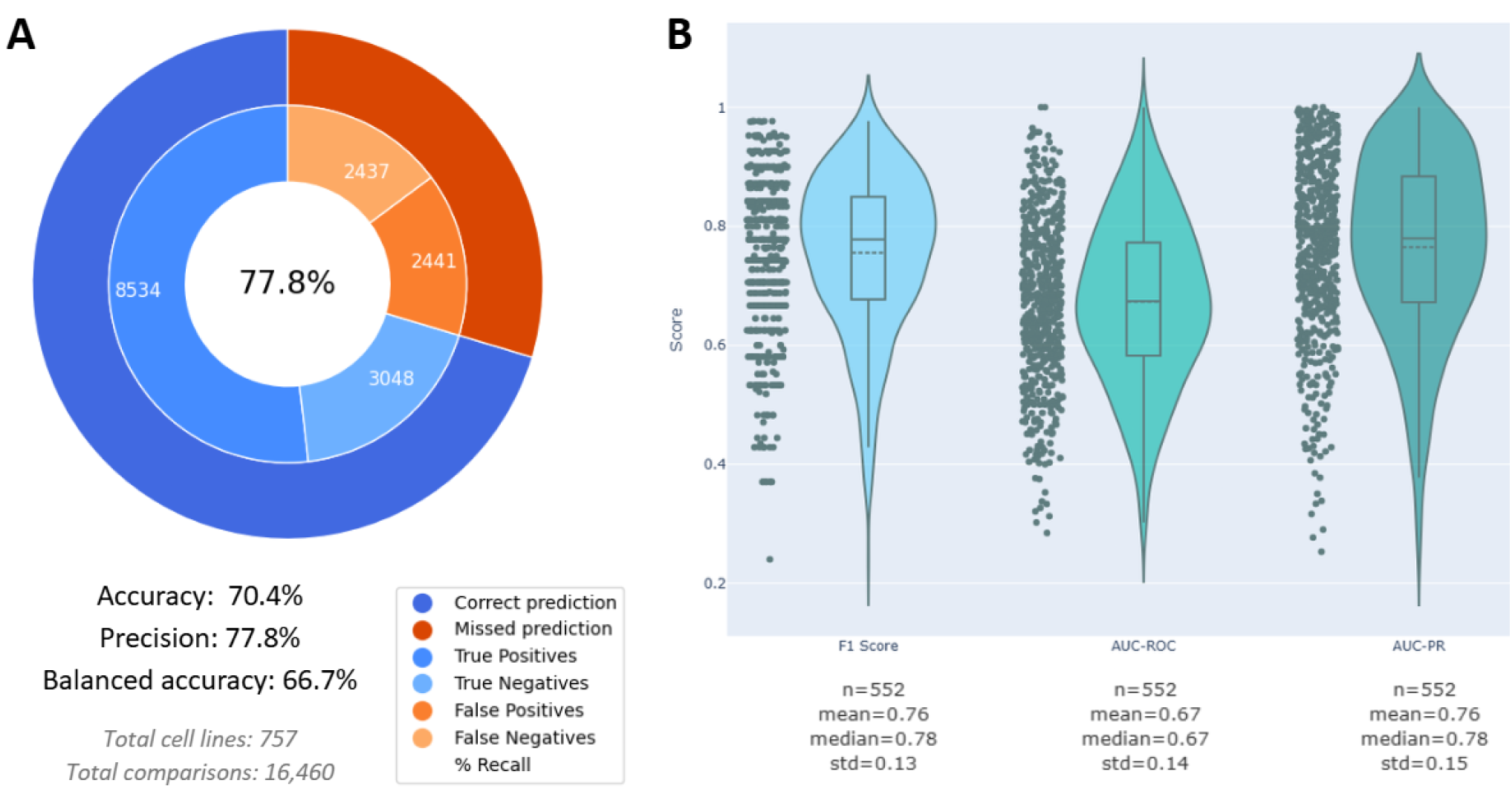
Predictive performance of calibrated in silico synergy predictions. **(A)** Ring plot showing mean accuracy, recall, and precision for predicting experimentally observed synergistic drug combinations across 16,460 cell line–combination comparisons in the Sanger-2024 dataset. **(B)** Violin plots showing the distribution of F1 score, ROC–AUC, and PR–AUC values across cell lines. Each point corresponds to a cell line metric. Centre lines indicate medians, boxes the interquartile range, and whiskers 1.5× the interquartile range.

Trafikk application to the independent Merck-2016 dataset yielded comparable classification accuracy (75.1%), supporting the general applicability of the pipeline. However, discrimination metrics could be computed for only a subset of cell lines (13 out of 38), due to limited class balance. Detailed performance metrics for Merck-2016 are provided in **Supplementary Fig S4**. Despite these constraints, overall classification performance remained directionally consistent with the primary analysis.

Together, these results indicate that the pipeline can *in silico* recreate experimental drug-combination responses, motivating further analyses of performance variability and mechanistic drivers. All subsequent analyses were performed using the Sanger-2024 dataset.

### Functional responses cluster primarily by drug combination rather than by cellular context

The nodes of the CFDv2 model were significantly enriched in 42 non-redundant KEGG pathways (**Supplementary Fig S5**). Pathway-level functional scores (of true positive and negative predictions) derived from these pathways were used for the downstream analyses reported below. These pathway-level scores are derived from the same Boolean simulations used to compute Bliss synergy excess from viability-related nodes serving as proxies for proliferation and apoptosis in Oris, and therefore represent a mechanistic decomposition of the perturbation effects underlying the synergy calculation (**Fig 1**).

Results of principal component analysis (PCA) revealed that the applied drug combinations constitute a major source of functional variability across cell lines, leading to a clear stratification in functional space (**Fig 2A**, **Supplementary Fig S6**). Unsupervised hierarchical clustering of pathway scores identified ten functional clusters with distinct response patterns (**Fig 2B**, **Fig 3**, **Supplementary Fig S1**). Clusters enriched in BCL2-targeting combinations (clusters 8-10) were characterized by the highest relative pathway scores for apoptosis and p53-associated programs consistently across cell lines, with quantitative variation between contexts (**Fig 2A**). A second group of clusters (clusters 2-7) showed response patterns dominated by one of the signalling axes targeted by the combination, including ERBB-centred (clusters 2-3), MEK-centred (cluster 4), and PI3K/AKT/mTOR-centred combinations (clusters 5-7), each displaying distinct relative modulation of immune-related, junctional, metabolic or cell-cycle-associated pathways.

**Fig 2.**
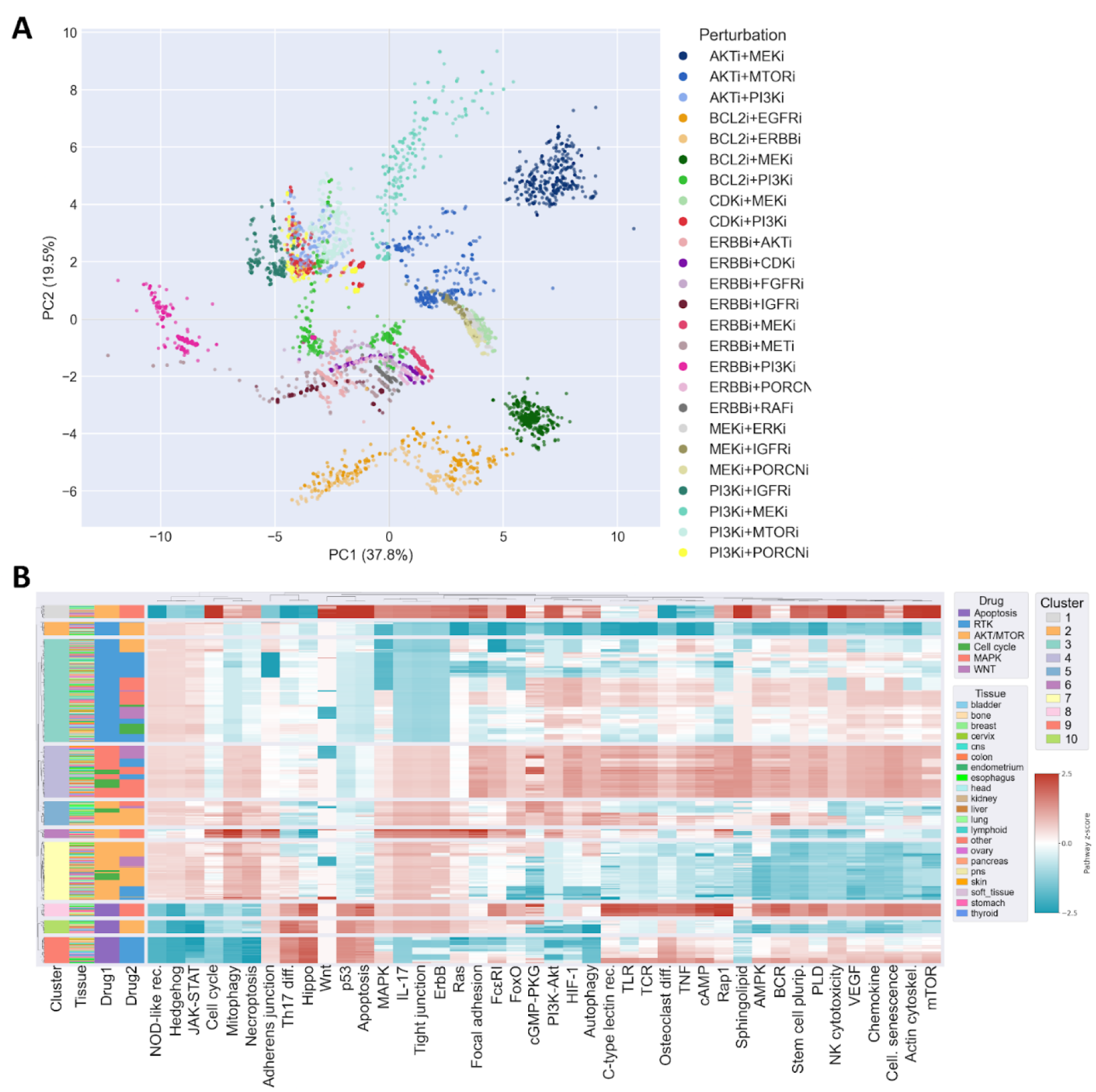
Global functional landscape of pathway responses to drug combinations. **(A)** Principal component analysis (PCA) of functional pathway scores, restricted to true positive and true negative responses. Each point represents a cell line treated with a given drug combination, and colours indicate the specific combination. **(B)** Heatmap of z-scored pathway scores for all combinations and cell lines, ordered according to the same hierarchical clustering (K = 1). Rows correspond to samples and columns to pathways.

**Fig 3.**
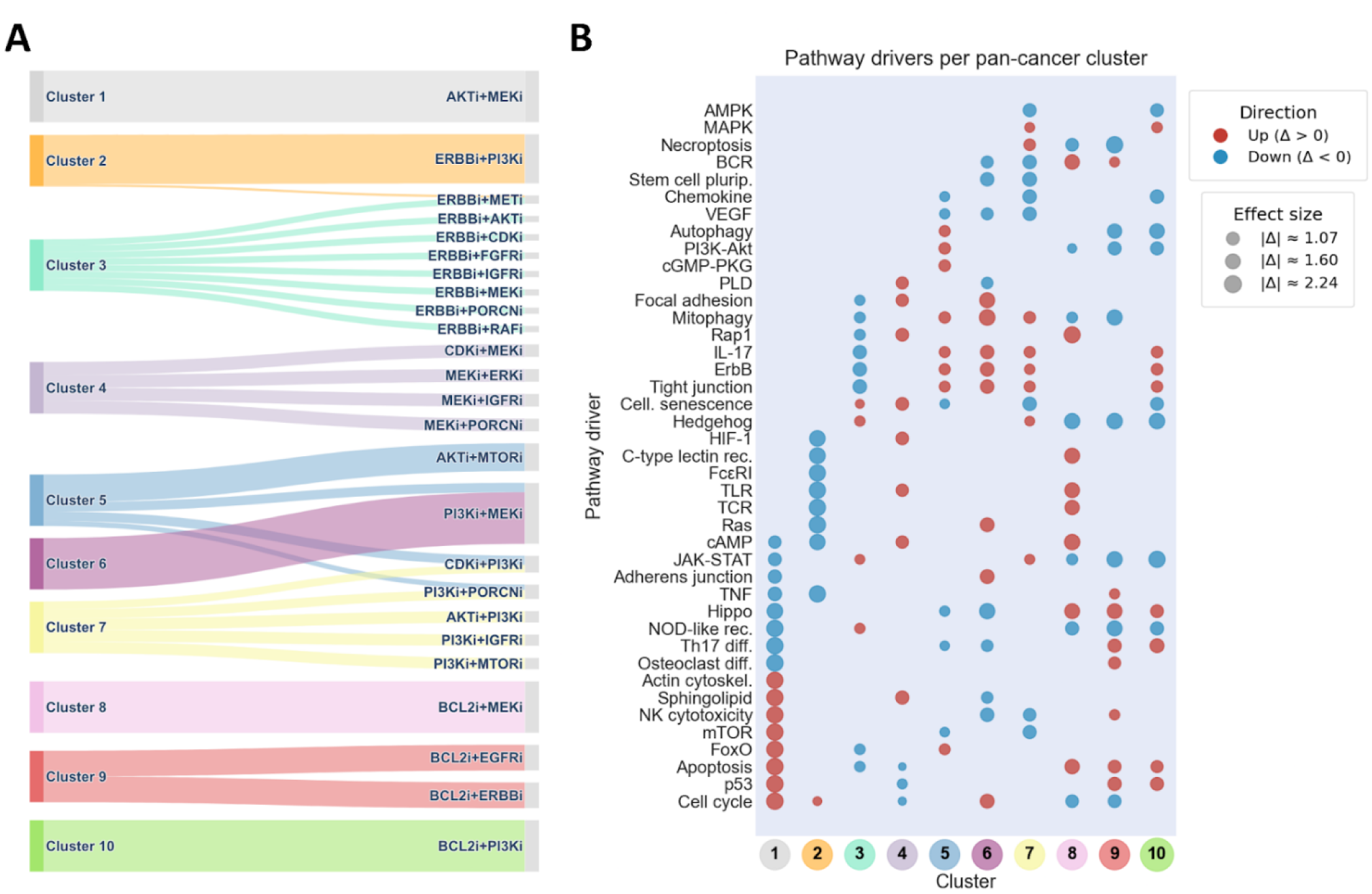
Cluster-specific functional programs underlying drug combination responses. **(A)** Sankey diagram linking drug combinations to functional clusters obtained by silhouette-optimized hierarchical clustering and pathway-level functional scores (K = 10). **(B)** Dot plot showing cluster-specific pathway divers. For each cluster, the top eight significantly enriched and depleted pathways (up and down) were identified by combining in-cluster versus out-of-cluster samples (Wilcoxon rank-sum test, FDR < 0.01) with a consistency threshold of ≥65%. Dot colour indicates the sign of the effect (Δ = μ_*in*_ ― μ_*out*_), and dot size reflects its absolute magnitude.

Markedly, AKT-MEK co-inhibition (cluster 1) was associated with a broader redistribution of pathway level scores than the other MEK-driven combinations (cluster 4), and AKT/PI3K/mTOR-driven combinations (clusters 5-7) (**Fig 2B**, **3**). Among the top significantly modulated pathways FDR < 0.01) were cell cycle- and stress-associated programs (*Cell cycle*, *p53*, *Apoptosis*, *mTOR*, *FoxO*), as well as immune- and cytoskeleton-related pathways (*Natural killer cytotoxicity*, *actin cytoskeleton*). In parallel, the most downweighted pathways included inflammatory and developmental signalling axes (*Th17 differentiation*, *TNF*, *Hippo*, *JAK-STAT*), consistent with a combined response pattern rather than a single dominant program. By comparison, MEK-driven combination responses displayed a more restricted functional profile. Downweighted pathways were largely limited to apoptosis (*Cell cycle*, *Apoptosis*, *p53*), whereas upweighted pathways primarily involved adaptive signalling programs (*HIF-1*, *cAMP*, *phospholipase D*). In contrast, AKT/PI3K/mTOR-driven combinations were characterized by prominent contributions from pathways closely associated with PI3K-AKT signalling and cell survival, as well as recurrent modulation of adhesion and immune-related pathways, consistent with a more axis-centred response. Overall, functional responses grouped primarily according to drug combination rather than cell line, with pathway modulation patterns reflecting the targeted signalling axis.

### Mechanistic pathway dynamics explain the AKTi and MEKi combination effect

To investigate how synergistic drug combinations translate into biological effects, we analysed the AKT inhibitor MK-2206 and the MEK inhibitor trametinib, a recurrent and robust combination across the cell line models. This pair was selected based on consistent pathway-level signatures across cancer types, indicating a combinatorial response independent from cancer type (**Fig 2A**, **3A**). Importantly, this combination was first identified as synergistic based on Bliss excess computed from viability-related output nodes, and the pathway-level analysis presented here dissects the signalling mechanisms underlying that quantitatively predicted effect.

In pathway-level analysis, AKT or MEK inhibition alone induced limited and partially overlapping pathway perturbations, whereas their combination produced a distinct reprogramming of signal propagation. In particular, this combination led to the activation of stress- and apoptosis-associated pathways, including p53 signalling, while attenuating survival and differentiation programs, which reveals non-additive pathway responses (**Fig 4A**).

**Fig 4.**
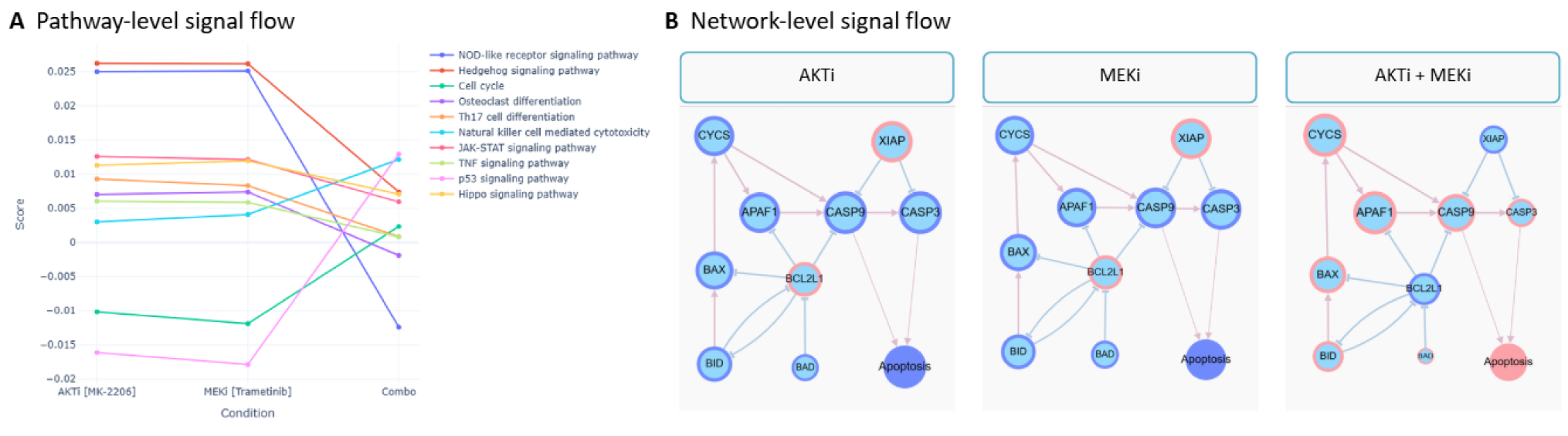
Pathway-level effects of a synergistic drug combination. **(A)** Pathway-level signal scores (precomputed from node-level data, Equation 4) are shown for AKTi, MEKi, and their combination. Pathways are ranked by an impact metric (Equation 5) that quantifies how much the combination differs in magnitude from the stronger single-drug effect. Median summarizes pathway-level scores across cancer types (equation 4). **(B)** Network-level signal propagation within the base network under the same conditions. A reduced visualization of the network is shown for pathway simplification purposes. Node scores across cancer types are overlaid, size reflects the absolute magnitude of the node score, and colour indicates the score sign, red for upregulated and blue for downregulated. The combination induces pathway and node-level changes not explained by single-drug effects, highlighting coordinated signal redistribution associated with synergy.

By mapping these effects onto the base network, we revealed that synergy originated from shifting the network balance, not from simply amplifying individual drug effects. While monotherapies maintained survival buffering, the combination of AKT and MEK inhibitors suppressed BCL2L1-mediated inhibition, triggered BAX activation, and initiated cytochrome-c-dependent CASP9-CASP3 signalling. This overcame XIAP-mediated restraint (**Fig 4B**). These results show that synergy occurs when combined perturbations break down survival buffering, pushing the cell toward an irreversible apoptotic state.

### TP53 status reveals distinct, context-dependent signalling routes converging on the same cellular fate

To investigate mechanistic sources of the functional heterogeneity within drug combinations, we first quantified pairwise differences in pathway-level responses across cell lines for each combination. Among all tested conditions, combined PI3K and BCL2 inhibition displayed the highest overall discordance between the cell lines (**Supplementary Fig S7**). We therefore selected this combination for in-depth analysis.

To uncover and exemplify the discordance, we used the most divergent cell line pair under combined PI3K and BCL2 inhibition: HO-1-u-1, derived from salivary gland carcinoma, and NUGC-4, derived from gastric cancer. Several of the pathways showing the largest differences between these cell lines were related to cell death and stress responses, including apoptosis, necroptosis, and autophagy-associated signalling (**Fig 5A**, upper panel). These pathways exhibited differences in both magnitude and direction between HO-1-u-1 and NUGC-4, indicating divergent downstream functional responses to the same perturbation.

**Fig 5.**
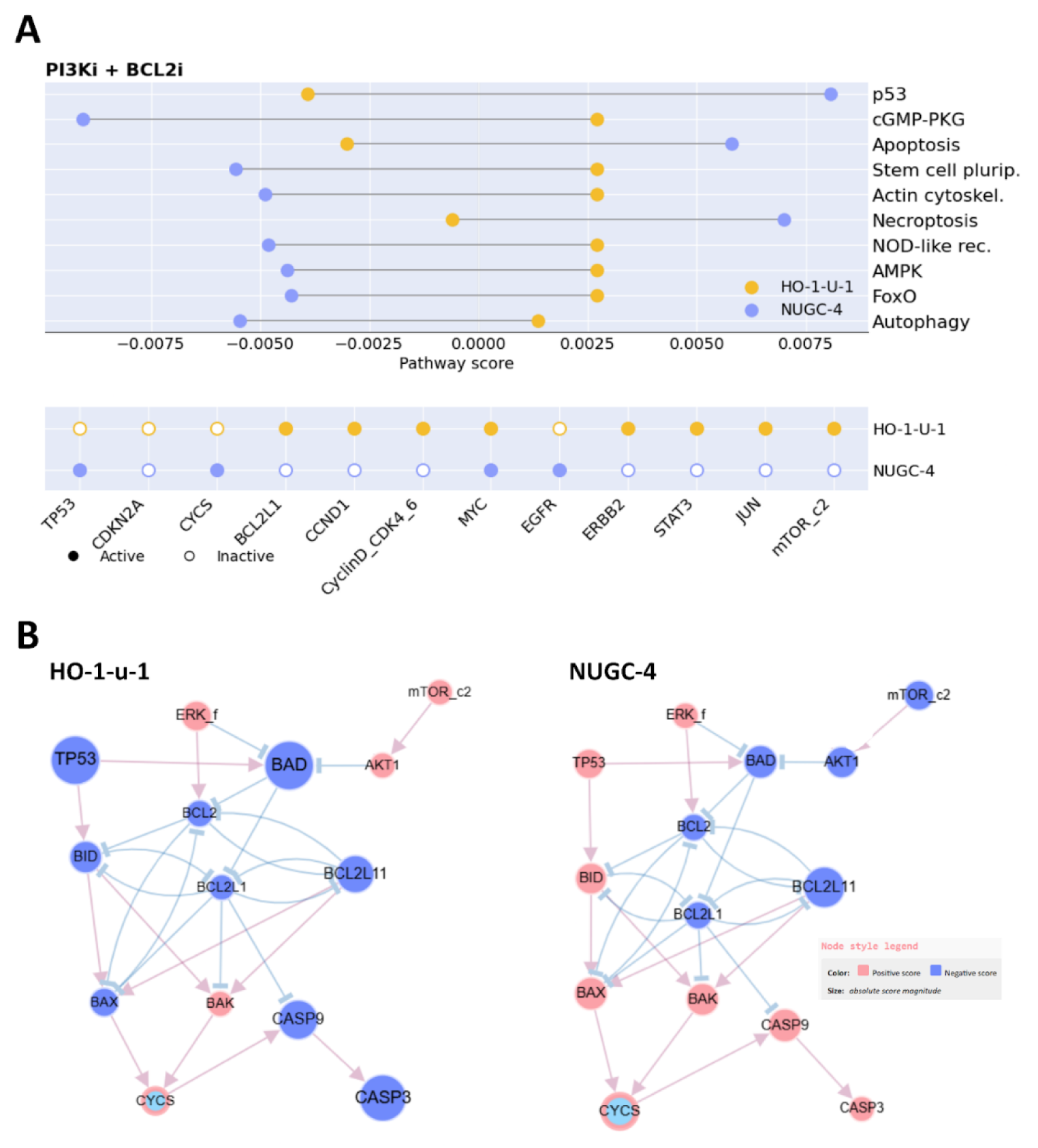
Cell-line-specific functional and apoptotic signalling differences under combined PI3K and BCL2 inhibition. **(A)** Pathway-level comparison of functional responses between HO-1-u-1 and NUGC-4 following combined PI3K and BCL2 inhibition. The dumbbell plot displays the 10 pathways showing the largest differences between cell lines, selected based on an absolute fold change greater than 1.5. Pathway scores quantify the signed bias of signal propagation toward each pathway in the Boolean models. The lower panel shows the baseline calibration states used to constrain model inference for each cell line. Filled circles indicate nodes constrained as active (1), and open circles indicate nodes constrained as inactive (0). **(B)** Cell-line-specific apoptotic signalling networks inferred from the same underlying network topology. Node colour represents the inferred activity state, and node size reflects the relative magnitude of node activity. Differences between HO-1-u-1 and NUGC-4 illustrate how identical perturbations lead to distinct patterns of signal propagation through the apoptotic subnetwork. Cytochrome c (CYCS) is highlighted for reference.

To contextualize these differences, we examined the baseline calibration states to constrain model inference (**Fig 5A**, lower panel). Notably, TP53 was constrained as inactive in HO-1-u-1 and active in NUGC-4, reflecting cell-line-specific baseline differences that are expected to influence downstream signalling responses.

We next examined how these differences were reflected at the network level by focusing on the apoptotic subnetworks (**Fig 5B**). Although both cell lines exhibited synergistic responses to PI3K and BCL2 co-inhibition, their inferred apoptotic signalling patterns differed substantially. In NUGC-4, multiple apoptotic components showed higher activity, consistent with a strong engagement of apoptotic signalling. In contrast, HO1-u-1 displayed limited activation of apoptotic nodes. Nonetheless, BAK and cytochrome c (CYCS) showed increased activity in HO-1-u-1, suggesting engagement of mitochondrial dysfunction despite reduced overall apoptotic signalling. Taken together, these results show that differences in baseline signalling states, exemplified by TP53 activity, can give rise to distinct downstream functional responses to identical drug combinations across cell lines.

### Trafikk provides a unified mechanistic and context-specific framework for drug synergy prediction

Trafikk builds upon a range of existing tools, pipelines, and methods for drug synergy prediction. Here, we position Trafikk within the current state of the art (**Table 1**, **Supplementary Table 4**).

**Table 1.**
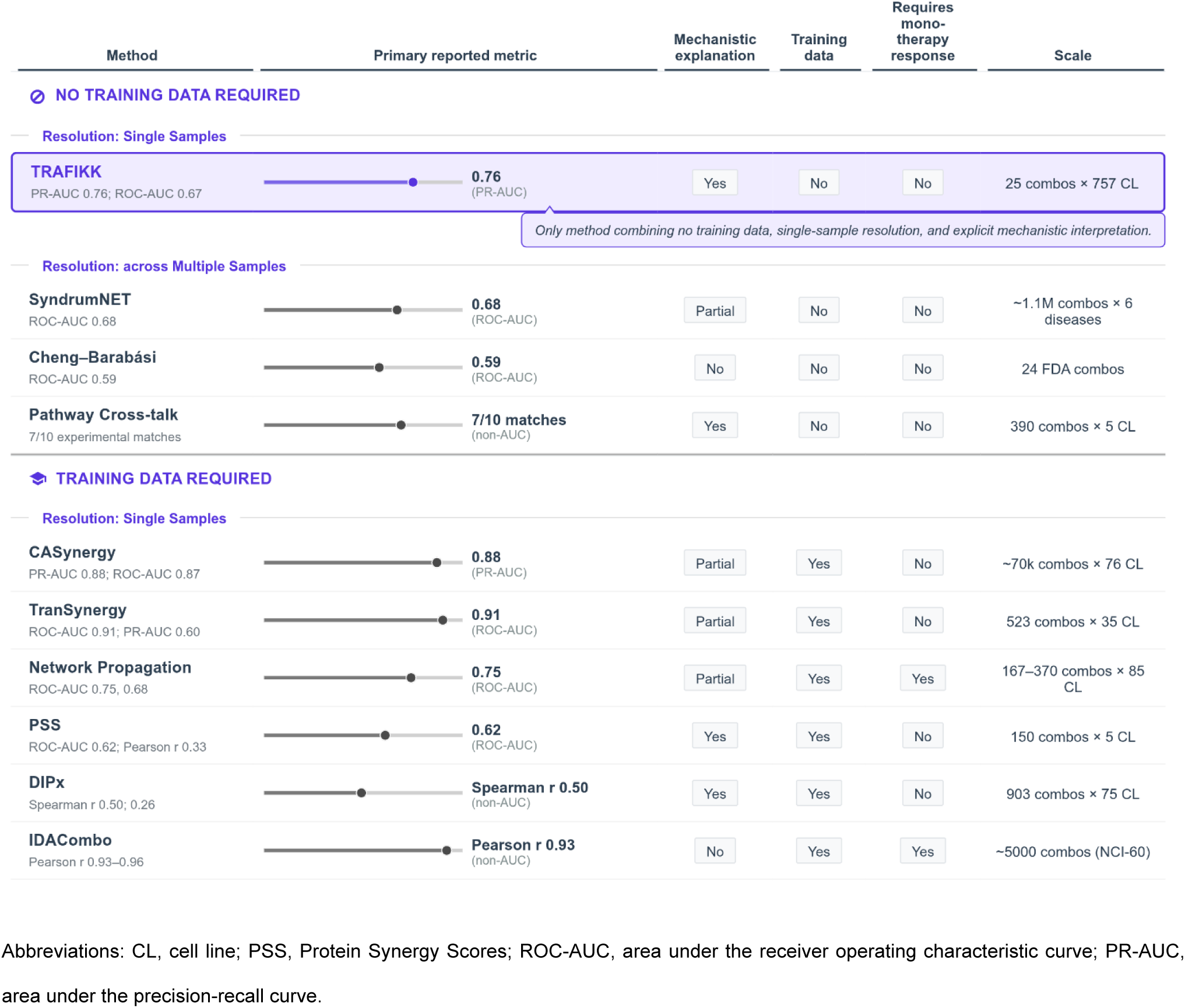
Trafikk in the landscape of computational drug synergy prediction methods.

| Method | Primary reported metric | Mechanistic explanation | Training data | Requires mono-therapy response | Scale |
| --- | --- | --- | --- | --- | --- |
| 🔓 NO TRAINING DATA REQUIRED |  |  |  |  |  |
| Resolution: Single Samples |  |  |  |  |  |
| **TRAFIKK**<br>PR-AUC 0.76; ROC-AUC 0.67 | **0.76**<br>(PR-AUC) | Yes | No | No | 25 combos × 757 CL |
| Only method combining no training data, single-sample resolution, and explicit mechanistic interpretation. |  |  |  |  |  |
| Resolution: across Multiple Samples |  |  |  |  |  |
| **SyndrumNET**<br>ROC-AUC 0.68 | **0.68**<br>(ROC-AUC) | Partial | No | No | ~1.1M combos × 6 diseases |
| **Cheng–Barabási**<br>ROC-AUC 0.59 | **0.59**<br>(ROC-AUC) | No | No | No | 24 FDA combos |
| **Pathway Cross-talk**<br>7/10 experimental matches | **7/10 matches**<br>(non-AUC) | Yes | No | No | 390 combos × 5 CL |
| 🔒 TRAINING DATA REQUIRED |  |  |  |  |  |
| Resolution: Single Samples |  |  |  |  |  |
| **CASynergy**<br>PR-AUC 0.88; ROC-AUC 0.87 | **0.88**<br>(PR-AUC) | Partial | Yes | No | ~70k combos × 76 CL |
| **TranSynergy**<br>ROC-AUC 0.91; PR-AUC 0.60 | **0.91**<br>(ROC-AUC) | Partial | Yes | No | 523 combos × 35 CL |
| **Network Propagation**<br>ROC-AUC 0.75, 0.68 | **0.75**<br>(ROC-AUC) | Partial | Yes | Yes | 167–370 combos × 85 CL |
| **PSS**<br>ROC-AUC 0.62; Pearson r 0.33 | **0.62**<br>(ROC-AUC) | Yes | Yes | No | 150 combos × 5 CL |
| **DIPx**<br>Spearman r 0.50; 0.26 | **Spearman r 0.50**<br>(non-AUC) | Yes | Yes | No | 903 combos × 75 CL |
| **IDACombo**<br>Pearson r 0.93–0.96 | **Pearson r 0.93**<br>(non-AUC) | No | Yes | Yes | ~5000 combos (NCI-60) |

Abbreviations: CL, cell line; PSS, Protein Synergy Scores; ROC-AUC, area under the receiver operating characteristic curve; PR-AUC, area under the precision-recall curve.

Several approaches, including TranSynergy (17) and CASynergy (8) integrate prior-knowledge networks, omics data, and drug-sensitivity measurements to train machine learning models that predict drug combinations, either as binary classifications or continuous scores (e.g. Loewe) (19). These methods typically rely on a training phase that optimizes predictive performance based on available data in a given condition (e.g. cancer type). This dependency on training data is also shared by other approaches, such as Network Propagation (20), IDACombo (21), DIPx (22), and Protein Synergy Scores (PSS) (**Table 3**) (18), the latter of which combines Boolean modelling with data-driven optimization and has shown good performance in breast cancer context. In contrast, a subset of network-based approaches, including the methods by Cheng-Barabási (23), SydrumNET (24), and Pathway Cross-talk (25), do not require training. These approaches leverage prior-knowledge networks and drug-target information to assess how drug combinations perturb disease-relevant regions of the network, typically through topological or proximity-based measures. While this makes them more readily applicable across conditions, their predictions are generally not tailored to specific cellular contexts.

Trafikk adopts a different strategy by integrating condition-specific omics data to generate cell line-specific Boolean models that serve as digital representations of the biological system. Once calibrated, these models can simulate arbitrary perturbations, including unseen drug combinations, without requiring additional data-driven training. Beyond prediction, Trafikk distinguishes itself through its mechanistic interpretability. While some other methods provide partial interpretability, such as pathway-level insights in DIPx or Pathway Cross-talk, protein-level contributions in Protein Synergy Scores (PSS), or post hoc explanations in deep learning approaches like TranSynergy and CASynergy (e.g. Shapley Additive Explanations (SHAP) or enrichment analyses), these are typically limited in scope or indirect through independent analyses. In contrast, Trafikk directly exposes the propagation of signals through the network, enabling coherent interpretation at the node, pathway, and network-module levels. This allows researchers not only to predict whether a combination is synergistic, but also to understand how and why the synergy emerges within a given biological context.

## Discussion

In this study, we present a mechanistic, network-based computational approach to systematically evaluate cancer drug combination responses under cell line-specific molecular contexts. The framework integrates cell-line-specific molecular context with drug-induced perturbations of signalling networks to model downstream cell-fate decisions related to proliferation and apoptosis. By using Bliss independence to quantify synergies, the pipeline enables direct comparison between simulated and experimental outcomes. Across two independent large-scale drug combination studies, this approach achieved consistent predictive performance, with both precision and recall above 77% for synergistic combinations. In addition, the proposed framework emphasizes mechanistic interpretability, allowing direct inspection of how drug effects propagate through signalling pathways. Together, these results demonstrate that network-based signal-propagation modelling can reproduce experimentally observed drug combination effects and provide mechanistic insight into the emergence of synergy.

To date, several computational approaches spanning machine learning and network-based models exists that leverage large-scale screening data and biological interaction information to identify synergistic combinations (6, 7, 27). However, only a few methods are available to support biological interpretation of combination synergies through feature attribution or pathway-level analysis (8). In the present study, in addition to predicting drug synergies, we analyse how single and combined drug perturbations reshape signalling activity. The motivation is two-fold; by focusing on pathway- and signal-propagation responses, the Trafikk framework first examines synergy as a result of combined drug perturbations and then evaluates it against experimental drug-screening data. This allows single and combination perturbations to be analysed and compared. Such a pathway perspective is increasingly relevant as there is a growing interest in higher-order drug combinations of more than two drugs, where experimental coverage remains limited and mechanistic insight can help guide exploration of the massive combinatorial search space.

Through the functional signal exploration enabled by the pipeline, we observed that drug combination identity was the primary source of variability in pathway-level response profiles, largely independent of the tissue origin of the treated cell lines. This is consistent with experimental observations from the large-scale drug screens used for validation, where several combinations induced relatively homogeneous viability responses across diverse cellular contexts(1, 2). Such behaviour is also expected given that the underlying network captures core, cancer-relevant signalling programs governing generic cell-fate decisions, rather than tissue-specific regulatory mechanisms. Interestingly, most combinations were functionally dominated by a single agent, indicating that one drug often dictates the signalling trajectory of the pair; such dominance effects have also been reported in molecular profiling studies and drug combination screens, and inferred from phospho-signalling datasets (3, 28). Notably, AKT-ERK dual inhibition, a recurrently reported synergistic combination (29, 30), deviated from this dominance pattern and produced a composite signalling signature. More generally, this example illustrates how synergistic drug responses can emerge when combined perturbations reorganize network signal flow in ways not accessible to single agents, highlighting the interpretive value of pathway- and signal-propagation analysis.

Beyond these global trends, our framework also revealed between-cell line heterogeneity for the same combination. Notably, PI3K-BCL2 co-inhibition produced synergistic effects in HO-1-u-1 and NUGC-4, yet with distinct signalling programs, illustrating that shared phenotypic outcomes do not require uniform execution mechanisms. Such divergence is consistent with the view that cell death pathways are not merely triggered but integrated within the molecular context of each cell line, including differences in p53 status and apoptotic priming. In this scenario, loss of functional p53 in HO-1-u-1 (31) may bias the response toward p53-independent, mitochondria-associated regulated cell death, in line with reports that p53 can redirect lethal stress toward mitochondrial respiration-dependent, apoptosis-dependent death programs (32). By contrast, NUGC-4, which retains wild-type p53 (33), exhibits broader engagement of canonical apoptotic components, and apoptosis can indeed be experimentally induced in this line via caspase-dependent mechanisms (33). Together, these observations indicate that phenotypic synergy can arise through distinct execution routes depending on apoptotic wiring, reinforcing that mechanistically similar drug synergies need not converge at the level of cell-death implementation.

As with other Boolean network-based approaches (34), the scope of Trafikk reflects a trade-off between network size and computational tractability. Recent methodological advances, such as the BooLEVARD framework developed by our group, enable quantitative analysis of signal propagation in Boolean models and facilitate this type of mechanistic analysis of perturbation effects (15). As network size and connectivity increase, the complexity of the attractor landscape grows rapidly, leading to a substantial rise in the cost of explicit signal-propagation analysis. For this reason, Boolean models often rely on abstractions that capture key signalling logic without explicitly representing all molecular details. In this context, generic or composite nodes may represent multiple biological instances, providing a practical way to balance biological expressiveness and computational feasibility while preserving interpretability.

Likewise, while the calibration procedure prioritizes models that admit stable states under baseline conditions, certain perturbations may shift model dynamics toward cyclic attractors (35), limiting the availability of steady-state readouts. Trafikk allows to partially mitigate this effect by increasing model ensemble size or adjusting sampling constraints, at the expense of additional computation. Finally, the use of Bliss independence provides a well-established (36) and conceptually compatible reference for synergy assessment within Boolean frameworks (10), while naturally abstracting away temporal dynamics, dose dependencies, and kinetic parameters that lie beyond the scope of logic-based modelling.

In conclusion, this study demonstrates that mechanistic network-based modelling can reproduce experimental drug combination responses while directly explaining how synergy emerges from drug-network interactions. By making pathway activity, signal propagation and cell-fate outcomes explicit, the framework offers a complementary alternative to purely predictive models. Looking ahead, combining such mechanistic representations with machine learning approaches could further improve both predictive performance and biological interpretability. In this way, Trafikk provides a practical foundation for interpreting large-scale drug screens and guiding the rational exploration of combination therapies.

## Methods

### Drug screen datasets

To evaluate the performance of the Trafikk pipeline, we used two independent experimental drug synergy datasets to compare our synergy predictions. The primary dataset was generated by the Wellcome Sanger Institute and thus named Sanger-2024 (2). The secondary dataset generated by Merck & Co., named Merck-2024 was mainly reserved for validation (1). Both datasets measured cell-based drug synergies for double perturbations under various cell line conditions (**Table 2**).

**Table 2.** Experimental drug synergy datasets.

| Experimental Dataset | Single Drugs | Combinations | Dose-combination matrix | Cell lines | Cancer Types |
| --- | --- | --- | --- | --- | --- |
| Sanger-2024 | 29 | 51 | 2 x 7 | 757 | 26 |
| Merck-2016 | 38 | 583 | 4 x 4 | 39 | 6 |

The Sanger-2024 dataset is a recently published large-scale drug screen that comprises 51 drug combinations tested across 757 cancer cell lines spanning 26 cancer types (2), and was used for downstream mechanistic and pathway-level analyses due to its scale and biological diversity. The Merck-2016 dataset is a large-scale drug screen that includes 538 drug combinations tested in 39 cancer cell lines across six cancer types (1), and was used as an independent benchmark to assess the generalizability of the pipeline.

Bliss excess for a drug combination was computed as defined in **Equation 1**:

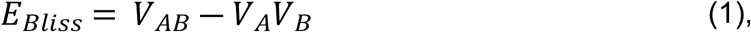

where *V*_*AB*_ denotes the viability under the combined perturbation, and *V*_*A*_ and *V*_*B*_ correspond to the respective single-perturbation conditions.

Bliss synergy scores were retained as continuous values for descriptive analysis. For performance evaluation, synergy identification was operationalized as a binary classification task. Drug-cell line pairs with ΔBliss < 0 were classified as synergistic, whereas ΔBliss ≥ 0 were classified as non-synergistic. In this viability-based formulation, negative values reflect deviation from Bliss additivity toward greater-than-expected reduction in viability, but do not necessarily imply a large or biologically significant synergistic effect. Importantly, the interpretation of numerical thresholds is scale-dependent and cannot be directly transferred from effect- or percentage-based Bliss implementations. Because the primary objective of this evaluation was directional classification rather than effect size stratification, the binary framework distinguishes only between deviation toward synergy (ΔBliss < 0) and non-synergistic responses, without assigning biological weight to marginal deviations from zero.

### Trafikk pipeline

Trafikk is a computational pipeline for *in silico* prediction of combinatorial drug responses using a network calibrated to specific cell-line contexts. The pipeline builds ensembles of logic-based network models, in which simulations of drug perturbations are performed to assess their impact on cancer signalling pathways. Trafikk is built upon existing software, including DrugLogics for generating the models (10), and BooLEVARD for computing the synergy scores and performing signal propagation analysis (15), and extends them with modules for data preparation, model calibration, and large-scale drug-screen analysis. The pipeline is organised into modules (**Fig 6**), which are outlined in the following sections. First, a network is calibrated in *Celios* with the biological context of each cell line, and an ensemble of models is generated in *Gitsbe* for each cell-line-network (**Supplementary Methods**). Then, the experimental drug combinations are translated into *in silico* combinations in *Drexpa* and evaluated in *Oris* to compute in silico synergy scores. The predicted synergy scores are compared with the experimental values in *Synco*, and *Siflex* is then used to investigate drug effects at the functional level and to generate mechanistic hypotheses. Full documentation is available at https://github.com/druglogics/trafikk.

**Fig 6.**
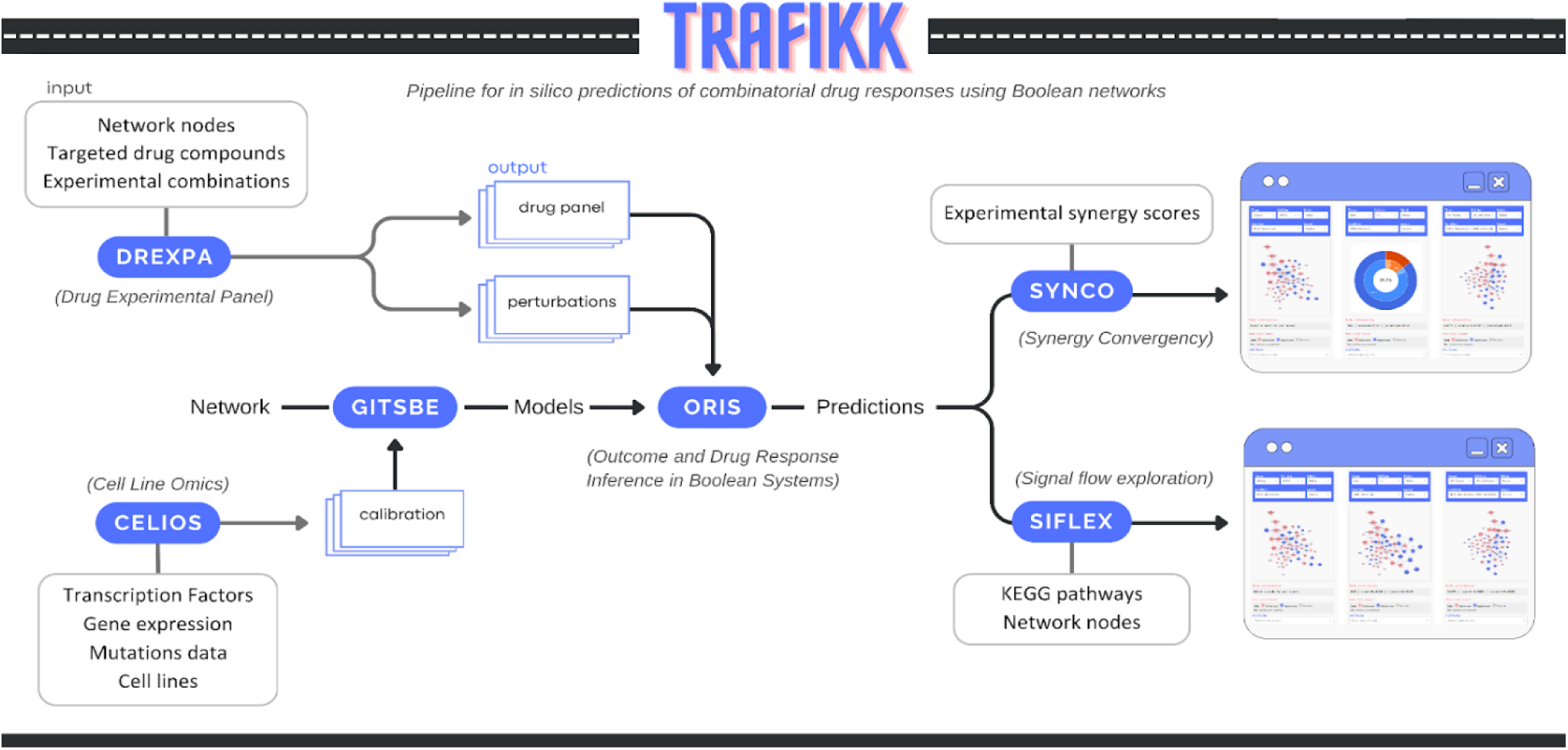
Overview and modular architecture of the Trafikk pipeline. The workflow illustrates the sequential integration of context-specific network calibration (Celios), ensemble model generation (Gitsbe), in silico drug translation (Drexpa), synergy scoring (Oris), predictive benchmarking (Synco), and functional analysis of drug effects.

#### Base network model

In the Trafikk pipeline, a base signalling network is required to initialise and optimise cell line-specific models. We used the Cell Fate Decision (CFD) model as the base network, as it captures core cancer-relevant pathways involved in cell survival, proliferation, and apoptosis in a tumour-agnostic manner.

To improve coverage of drug targets present in the Sanger-2024 datasets, we manually expanded the network (CFDv2). The original model already covered a subset of drug targets, and the expansion was limited to three drug target nodes and seven supporting interactions. Each node is annotated with the HGNC protein symbol(s), and each edge includes a PubMed identifier (PMID) for validation. Full model details are provided in **Supplementary Tables S1–S2**.

#### Celios module

To personalise the base network with cell-line-specific parameters, we first collected and integrated omics information using the Celios (Cell Line Omics) module to define node activities. Using the entities represented in the CFDv2 model, the module integrates gene mutation, gene expression, and transcription factor activity information, specific to the selected cell lines, with the generic network (**Supplementary Methods**). We limited the analysis to binarized mutations, CNV (GISTIC +2 coded as 1 and GISTIC -2 coded as 0) (Mermel, et al., 2011), and transcription factor activity as input to generate the activity matrix (**Supplementary Table S3**). The output is a calibration file for each cell line that will later be used in the next modules. Celios also adds a condition to the calibration file to select models in a *proliferation*-like state in the Gitsbe module, requiring the normalized *globaloutput* to be equal to 1. Here, the *globaloutput* represents the weighted aggregate activity of the module outputs, rescaled to the 0,1 interval (**Supplementary Methods**).

#### Drexpa module

To represent the drug panel and combinations from the experimental drug screen datasets in the Trafikk pipeline, we developed the Drexpa (Drug Experimental Panel) module to automatically generate pipeline drug profiles (PD_profiles) for each drug and the respective combinations. Drexpa integrates information from multiple public resources, including GDSC, OpenTargets Platform, UniProt, BindingDB, and ChEMBL, to query protein targets associated with each drug in the experimental dataset (**Supplementary Methods**). Each protein target is mapped to the CFDv2 model nodes, and drugs without a mapped target are excluded from the analysis. Therefore, we focused on a portion of experimental datasets (**Table 3**). With the drug panel, Drexpa generates a perturbation file for each cell line. Depending on the format of the drug screen dataset, it can be a generic perturbation file or specific to each cell line (**Supplementary Methods**).

**Table 3.** In silico drug screening data.

| Pipeline Dataset | Single drugs | Unique target profiles* | Cell lines | Cancer Types | Combinations |
| --- | --- | --- | --- | --- | --- |
| Sanger-2024 | 17 | 15 | 757 | 26 | 25 |
| Merck-2016 | 7 | 6 | 38** | 6 | 21 |

\* Unique target profiles correspond to single protein targets or a set of protein targets associated with an experimental drug.
\*\* The screen included the ovarian cancer cell line UWB1.289 and its BRCA1-reconstituted derivative (UWB1.289+BRCA1). However, no distinct SIDM was available for the BRCA1-reconstituted derivative and therefore the derivative line was excluded from the analyses.

#### Oris module

Oris (Outcome and Drug Response Inference in Boolean Systems) is a module built on top of the recently described BooLEVARD (15) for estimating drug combination effects from ensembles of Boolean models. Oris operates on model ensembles previously calibrated to experimental data and quantifies the impact of single and combined perturbations on downstream signalling, providing a mechanistic proxy for drug sensitivity at the network level.

To reduce the computational burden on large models or large perturbation sets, Oris implements a sampling step. Models are randomly selected and their performance is evaluated using BooLEVARD’s CountPaths method (15); models that complete within a user-defined threshold are retained. Sampling continues until the requested number of models is reached or no more valid models remain.

For synergy quantification, a subset of output nodes was designed to represent proliferation and apoptosis (**Supplementary Methods**). For each model instance and perturbation condition, Oris computes node-level path-count scores and, when multiple stable states satisfy imposed constraints (e.g. media conditions or other calibration data), averages them to obtain a single model-level response.

A viability score is defined with smooth normalisation, as defined in **Equation 2**:

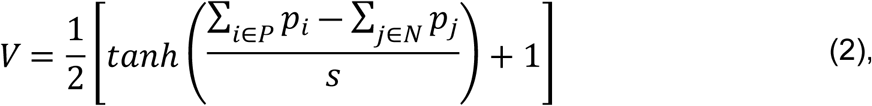

where *p*_*k*_ denotes the signed path-count score of output node *k*, *P* denotes pro-proliferative nodes, *N* denotes antiproliferative nodes, and *s* was calibrated from the empirical distribution of the raw viability signal to avoid saturation of the hyperbolic tangent mapping. Viability scores (**Equation 2**) were subsequently capped at the corresponding unperturbed value and normalised by the control, such that the unperturbed condition attained a value of *V* = 1.

In addition to computing synergy scores, Oris can quantify signed signal transduction paths toward all nodes in the network, enabling mechanistic inspection of how perturbations affect signal propagation signalling across the model (**Supplementary Methods**).

#### Synco module

To automatically evaluate all pipeline predictions against experimental datasets, we used the Synco (Synergy Comparison) module from DrugLogics, which integrates and compares *in silico* predictions with experimental synergy scores (**Supplementary Methods**). The module was adapted to integrate Oris synergy output and harmonise both *in silico* and experimental synergy data, enabling the calculation of standard decision-analytic metrics, such as accuracy, precision, and recall, as well as standard classification metrics, including the Area Under the Receiver Operating Characteristic Curve (AUC-ROC), the Area Under the Precision-Recall Curve (AUC-PR), and the F1 Score, and summarised using different plots (https://github.com/ViviamSB/SYNCO).

#### Siflex module

In addition to synergy data, the Oris module also generates node-level path-count scores, which can be used to interpret the activation and inhibition of signalling pathways inside the base model. Therefore, to investigate the synergistic drug effects at the functional level and generate mechanistic hypotheses, we developed the Siflex (Signal Flow Exploration) module, which focuses on explaining where and how these synergistic effects manifest. The module processes Oris outputs alongside the base network and KEGG pathway annotations.

First, to interpret node-level signalling information at a functional level, a pathway-based analysis of single and double-perturbation outcomes is performed by integrating BooLEVARD-derived path-count scores (15) into pathway-level activity measures. Nodes in the network were annotated to pathways using the KEGG database (26). Pathway enrichment is assessed using a hypergeometric test with Benjamini-Hochberg correction (FDR ≤ 0.01), and only significant enriched pathways were retained for downstream analyses. In addition, pathway overlap was quantified using the Szymkiewicz-Simpson coefficient.

For each model instance, node-level signed path-count scores are averaged across the selected stable states, yielding a single value *p*_*i*_ per node. These values were transformed into normalised node-level scores according to **Equation 3**:

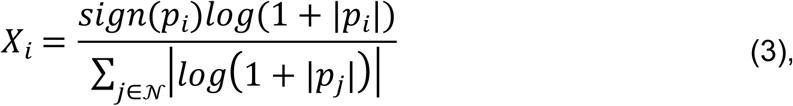

where *N* denotes the set of all nodes in the model. This transformation applies signed logarithmic scaling to attenuate large-magnitude path-count values while preserving directionality, followed by normalisation by the total absolute signal across the model.

Pathway-level scores were then computed by aggregating node-level scores for each pathway according to **Equation 4**:

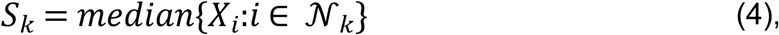

where *N*_*k*_ denotes the set of nodes annotated to pathway k. The median was used to obtain a robust summary of pathway-level activity.

These pathway-level scores are then used as input to support systematic comparison and visualization of single-drug and combination responses in Siflex. For combinations previously classified as synergistic using Bliss independence, Siflex computes a pathway impact score that quantifies the additional pathway perturbation induced by the combination relative to either single drug. For a given pathway k, the impact score is defined as **Equation 5**:

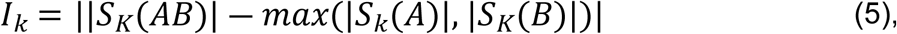

where *S*_*k*_ denotes the pathway-level scores under drug *A*, drug *B*, and the combination *AB*. This score is used to rank pathways and select subsets for downstream network- and pathway-level exploration (**Supplementary Methods**), without redefining or re-estimating drug synergy.

Together, this framework provides a consistent link between node-level signal propagation, pathway-level summarization, and downstream exploration of synergistic drug effects. By integrating normalized pathway scores and the pathway impact metrics within a modular analysis pipeline, Siflex enables systematic comparison of single-drug and combination responses across biological contexts. Detailed descriptions of data processing steps, visualization options, and interactive exploration are provided in **Supplementary Methods**.

### Unsupervised analysis of pathway-level functional responses

Pathway-level functional scores (**Equation 4**) were used as input for unsupervised analyses aimed at characterizing global response patterns across drug combinations and cellular contexts. Prior to multivariate analyses, feature matrices were standardized across samples by centring each pathway score to zero mean and scaling to unit variance (z-scores). False positives and false negatives were not used in this analysis.

Principal component analysis (PCA) was performed on the standardized pathway-level functional scores to identify dominant sources of variability across double-drug perturbations. Projections were computed using singular value decompositions and visualized in two dimensions. To identify recurrent functional response patterns, unsupervised hierarchical clustering was applied to the pathway-level functional scores. Hierarchical clustering was performed using average linkage, and the optimal number of clusters was determined by silhouette optimization, resulting in K = 10 functional clusters (**Supplementary Fig S1**).

Cluster assignments were used for downstream analyses and visualization. To identify cluster-specific pathway drivers, pathway scores within each cluster were compared against all remaining samples using a Wilcoxon rank-sum test. P values were adjusted for multiple testing using the Benjamini-Hochberg procedure, and pathways were considered cluster-specific if they met an FDR threshold of < 0.01 and showed consistent enrichment or depletion in at least 65% of cluster members.

In addition, cosine distances between pathway-level functional signatures were computed pairwise across cell lines within each drug combination to assess intra-combination functional variability. These distances were used to quantify the consistency of functional responses across cellular contexts for a given combination.

### Use of Generative Artificial Intelligence (AI)

During manuscript preparation, the authors used ChatGPT (OpenAI, GPT-4.5) and Grammarly for language refinement, structural editing, and improvement of clarity and readability. The tools were used to support phrasing and conciseness. The graphical abstract and Fig 6 were solely created by the authors using Canva, with no third-party copyrighted material. Table 3 was generated with assistance from the AI tool Claude (Anthropic). The authors reviewed and validated all outputs. No generative AI tools were used for data analysis, result interpretation, or the generation of original scientific content. All conceptual development, methodological design, analysis, and final editorial decisions were performed by the authors, who take full responsibility for the content of the manuscript.

## Data availability

Trafikk’s source code, installation instructions, and usage tutorial are freely available at https://github.com/druglogics/trafikk.

## Author Contributions

M.F. and V.S.B. contributed equally to this work in conceptualisation, investigation, formal analysis, data curation, writing (original draft), and visualisation. V.S.B developed the code for the Celios, Drexpa, and Synco modules. Kr.L. developed the code for the DrugProfiler database tool used in the Drexpa module. M.F., with support from J.Z. in conceptualization, developed the code for the Oris module. V.S.B and M.F. developed the code for the Siflex module. E.T., J.Z., T.A., Ka.L., and Å.F. contributed to supervision and writing (review & editing). T.A., Ka.L. and Å.F were responsible for project administration and funding acquisition.

## Supporting information

**Supplementary Fig S1.** Silhouette optimization for hierarchical clustering of pathway-level functional scores.

**Supplementary Fig S2.** Predictive performance across 26 cancer types in the Vis-2024 dataset. Ring plots summarizing classification performance for each of the 26 cancer types in the Vis-2024 dataset. For each cancer type, outer rings represent correct versus missed predictions, and inner rings show true positives, true negatives, false positives, and false negatives. Accuracy, precision, and balanced accuracy are reported below each plot, along with the number of evaluated cell lines per tissue, and recall is reported in the center of each ring.

**Supplementary Fig S3.** Distribution of performance metrics across cancer types in the Vis-2024 dataset. **(A)** Heatmap showing F1 score, ROC–AUC, and PR–AUC values per cancer type. **(B)** Violin plots displaying the distribution of ROC–AUC scores across selected tissues. The dashed horizontal line indicates the random classification threshold (ROC-AUC = 0.5). The accompanying table summarizes per-tissue statistics, including mean, median, minimum, maximum, standard deviation, and the number of cell lines with ROC-AUC > 0.5.

**Supplementary Fig S4.** Predictive performance in the O’Neil-2016 dataset. **(A)** Ring plot summarizing overall classification performance across cell line-combination comparisons in the O’Neil-2016 dataset. **(B)** Box plots showing F1 score, ROC–AUC, and PR–AUC distributions across evaluable cell lines. Metrics were computed only for cell lines containing both synergistic and non-synergistic experimental combinations. (C) Heatmap displaying per–cell line performance metrics (F1 score, ROC–AUC, PR–AUC).

**Supplementary Fig S5.** Overlap coefficient matrix between KEGG pathways significantly enriched for the model nodes (FDR < 0.01). For each pathway pair, the Szymkiewicz-Simpson overlap coefficient was computed based on gene sets restricted to the model foreground. Pathways were hierarchically clustered (average linkage, Euclidean distance) and only the lower triangle is displayed for clarity.

**Supplementary Fig S6.** Principal component analysis (PCA) of pathway-level functional scores derived from BooLEVARD. Each point represents a cell line under a given double-perturbation condition, restricted to true positive and true negative responses. Faceted panels show individual combinations with points colored by tissue of origin.

**Supplementary Fig S7.** Inter-cell line heterogeneity across drug combinations. Scatterplots show pairwise cosine distances between cell-line functional pathway signatures for each drug combination. Each point represents a pair of cell lines treated with the same combination. Combinations are ordered along the y-axis based on median cosine distances, ranging from more broadly heterogeneous patterns to more consistent responses, and points are colored according to the functional cluster assignment.

**Supplementary Table S1.** Node annotations for the Cell Fate Decision (CFDv2) network.

**Supplementary Table S2.** Edge annotations for the Cell Fate Decision (CFDv2) network, including interaction types and regulatory directionality.

**Supplementary Table S3.** Calibration data used to generate cell line-specific Boolean models.

**Supplementary Table S4.** Trafikk in the landscape of computational drug synergy methods. Calibration data used to generate cell line-specific Boolean models. Abbreviations: CL, cell line; PPI, protein-protein interaction; CNV, copy number variation; TF, transcription factor; HSA, Highest Single Agent; IDA, Independent Drug Action; PCI, pathway cross-talk index; PSS, Protein Synergy Scores; ROC-AUC, area under the receiver operating characteristic curve; PR-AUC, area under the precision-recall curve; SHAP, Shapley Additive Explanations.

